# Evidence for a competitive relationship between executive functions and statistical learning

**DOI:** 10.1101/2023.01.19.524710

**Authors:** Felipe Pedraza, Bence C. Farkas, Teodóra Vékony, Frederic Haesebaert, Romane Phelipon, Imola Mihalecz, Karolina Janacsek, Royce Anders, Barbara Tillmann, Gaën Plancher, Dezso Nemeth

## Abstract

The ability of the brain to extract patterns from the environment and predict future events, known as statistical learning, has been proposed to interact in a competitive manner with prefrontal lobe related networks and their characteristic cognitive or executive functions. However, it remains unclear whether these cognitive functions also show competitive relationship with implicit statistical learning across individuals and at the level of latent executive function components. In order to address this currently unknown aspect, we investigated, in two independent experiments (N_Study1_ = 186, N_Study2_ = 157), the relationship between implicit statistical learning, measured by the Alternating Serial Reaction Time task, and executive functions, measured by multiple neuropsychological tests. In both studies, a modest, but consistent negative correlation between implicit statistical learning and most executive function measures was observed. Factor analysis further revealed that a factor representing the verbal fluency and complex working memory seemed to drive these negative correlations. Thus, an antagonism between implicit statistical learning and executive functions might specifically be mediated by updating component of executive functions or/and long-term memory access.

## Introduction

Statistical learning (SL) is a fundamental function of human cognition that allows the implicit extraction of probabilistic regularities from the environment, even without intention, feedback, and reward, and is crucial for predictive processing (Aslin, 2017; Conway, 2020; Kaufman et al., 2010). SL contributes to the acquisition of language (Ullman et al., 2020), motor (Hallgató et al., 2013; Verburgh et al., 2016), musical (Rohrmeier & Rebuschat, 2012; Romano Bergstrom et al., 2012; Shook et al., 2013) and social skills (Baldwin et al., 2008; Parks et al., 2020; Ruffman et al., 2012), as well as habits (Graybiel, 2008; Horváth et al., 2022; Szegedi-Hallgató et al., 2017). SL can occur incidentally, without awareness and the intention to learn (Destrebecqz & Cleeremans, 2001; Fu et al., 2010; Kóbor et al., 2017; Song et al., 2007a, 2007b; Vékony et al., 2022). SL does not function in an isolated manner, but in either cooperative or competitive interactions with other cognitive processes (Poldrack & Packard, 2003, Conway, 2020). Here, we aim to investigate the interaction between implicit SL and executive functions, and to determine which aspects of executive functions show a positive (cooperative) and which a negative (competitive) relationship with statistical learning.

The competition hypothesis was coined in the framework of interactive memory systems (Batterink et al., 2019; Freedberg et al., 2020; Janacsek & Nemeth, 2022; Poldrack & Packard, 2003). According to this framework, learning can rely on either the basal ganglia-based procedural system, or the medial temporal lobe-based declarative system. Larger reliance on one implies a smaller reliance on the other. Initial evidence for this hypothesis has come both from animal (Eichenbaum et al., 1988; Packard et al., 1989) and human neuroimaging studies (Poldrack et al., 1999, 2001). Later, studies rooted in computational neuroscience and reinforcement learning also shed more light on the role of models in guiding learning in these different systems (Daw et al., 2005). It was proposed that the distinction between the declarative and procedural systems might map onto the distinction between model-based and model-free learning algorithms (Doll et al., 2015; Shohamy & Daw, 2014). Model-based learning processes build and make use of a model of the environment to flexibly guide choice, but are more computationally demanding. In contrast, model-free learning processes are computationally less expensive as they rely only on recent outcomes; however, this simplicity allows for less flexibility and sensitivity to contingency changes. The brain is assumed to arbitrate between these two learning systems in a dynamic fashion both during the completion of individual tasks (Doyon et al., 2009; Lee et al., 2014; Poldrack et al., 2001), and also throughout the lifespan (Decker et al., 2016; Nemeth, Janacsek, & Fiser, 2013; Palminteri et al., 2016). Multiple possible arbitration mechanisms have been described, including direct neuroanatomical connections between basal ganglia and medial temporal cortex, differential effects of neuromodulators, and indirect cross-inhibition via prefrontal control processes (Freedberg et al., 2020). The present study focuses on this last mechanism.

The top-down cognitive processes necessary for the flexible cognitive control of behaviour are often collectively referred to as executive functions (EF) (Diamond, 2013; Koechlin, 2016; Miyake & Friedman, 2012). These processes are usually required when we encounter novel, unusual, or constraining situations. They include cognitive functions such as attentional control, cognitive flexibility, cognitive inhibition and working memory updating (Diamond, 2013; Miyake et al., 2000). Studies exploring the neural basis of EF consistently show that these cognitive processes rely heavily on the prefrontal cortex (PFC) (Badre et al., 2010; Braver et al., 2003; Koechlin et al., 2003; Miller & Cohen, 2001). Moreover, individual differences in EF ability have been associated with individual differences in local prefrontal neural activity (Osaka et al., 2003), as well as functional connectivity between PFC and other brain regions (Kondo et al., 2004). Here, we focus on the question of whether and how individual differences in executive functions might affect statistical learning.

Prefrontal EF has been implicated in arbitrating between learning systems by multiple studies. Lee et al. (2014), for example, showed that lateral PFC and frontopolar cortex seem to encode the reliability associated with both model-based and model-free learning systems, as well as the output of the arbitration process. Importantly, their functional connectivity results also suggested that the arbitration mechanism might work primarily by suppressing model-free learning, when it deems model-based learning to be more beneficial. An intriguing possibility is that the procedural, model-free system is the ‘default’ learner that is overridden by PFC control involvement. This hypothesis seems to be in line with a series of results that show a negative relationship between statistical learning and prefrontal lobe control processes at both the behavioural and the neural level. For instance, disruption of PFC function by transcranial magnetic stimulation (Ambrus et al., 2020; Smalle et al., 2017), by hypnosis (Nemeth, Janacsek, Polner, et al., 2013), or by cognitive fatigue (Borragán et al., 2016) and engagement of prefrontal lobe resources using dual task conditions (Filoteo et al., 2010; Smalle et al., 2022) all have been reported to increase statistical learning performance. Moreover, multiple neuroimaging studies have also revealed that statistical learning seems to be associated with generally decreased functional connectivity both within PFC circuits and between the PFC and other networks (Park et al., 2022; Tóth et al., 2017). Our goal is to test whether weaker PFC-dependent executive functions could lead to better statistical learning across individuals. That is, do people with relatively weaker executive functions have relatively better statistical learning ability?

The little interindividual differences research carried out so far has suggested that PFC-dependent cognitive functions might relate negatively to statistical learning capability. For instance, higher EF ability has been found to be negatively related to SL ability across individuals (Virag et al., 2015). However, several studies have indicated that working memory is independent from statistical learning (reviewed in Janacsek & Nemeth, 2013). Furthermore, some recent studies have found positive associations between statistical learning and EF ability (Park et al., 2020; Petok et al., 2022). Therefore, the relationship between SL and PFC-supported cognitive functions needs further examination in order to disentangle the still somewhat puzzling relation between the two neurocognitive mechanisms (Janacsek & Németh, 2015). The relationship between implicit statistical learning and PFC-supported cognitive functions has been empirically explored in two studies (Park et al. 2020; Virag et al., 2015); however, both studies had relatively low sample sizes (22 and 40, respectively) leading to low statistical power (Button et al., 2013), and inability to establish reliable correlations between the tasks (Schönbrodt & Perugini, 2013). As neither study included a replication sample, the robustness of their results is also currently unknown. Furthermore, they only studied the relationship between EF and SL at the task level, without considering the possibility that a pattern might instead emerge at the level of latent EF components, tapped into by multiple tasks. Such latent variable approaches might also lead to relatively higher reliability estimates of cognitive abilities, which is extremely important in inter-individual differences research (Haines et al., 2023). However, the latent structure of EF abilities is itself a contentious issue, with a recent large-scale analysis failing to find measurement models that consistently showed a good fit (Karr et al., 2018).

In this study, we aimed to investigate the relationship between EF and SL, using two large, independent samples acquired in two different studies, offering an internal replication and a far larger overall sample size than previous studies. Both studies measured implicit SL ability using the alternating serial reaction time task (ASRT), a valid and reliable task of implicit SL (Buffington et al., 2021; Farkas et al., 2023), and EF ability using a wide variety of well-characterized neuropsychological tasks. Reasoning in terms of competitive neurocognitive systems, we hypothesized that a negative correlation would be found between EF ability and SL performance. However, due to the heterogeneous findings in the literature regarding SL – EF relationships and the structure of EF itself, we did not formulate strong hypotheses regarding specific tasks and latent EF components. Instead, we focused on assessing EF – SL relationships in a data-driven manner, at both the level of individual EF tasks, as well as the level of latent EF components, tapped into by multiple tasks assessing EF, extracted by exploratory factor analysis.

## Methods

### Participants

For Study 1, conducted in France, participants took part in a two-day experiment. One hundred eighty nine healthy young adults were included with the following criteria: participants were right-handed, aged under 35 years, with musical training (as measured by practice) inferior to ten years, declared not having active neurological or psychiatric conditions, and declared not be taking any psychoactive medication. Among the 189 participants, two did not come back for the second session and one did not comply with the task instructions on the first session. Thus, the data from the remaining 186 subjects is presented in this study. All participants provided signed informed consent agreements and received financial compensation for their participation. The relevant institutional review board (i.e., the “Comité de Protection des Personnes, CPP Est I” ID: RCB 2019-A02510-57) gave ethical approval for the study.

In Study 2, conducted in Hungary, 180 participants took part in a multi-session experiment. Criteria were the same as in Study 1, with the exception of being under 35 and having no musical expertise. From this pool, we excluded 23 subjects who had missing data on any of the EF tasks. This study was approved by the United Ethical Review Committee for Research in Psychology (EPKEB) in Hungary (Approval number: 30/2012) and by the research ethics committee of Eötvös Loránd University, Budapest, Hungary. Descriptive statistics of both samples are presented in Table 1.

**Table 1.**
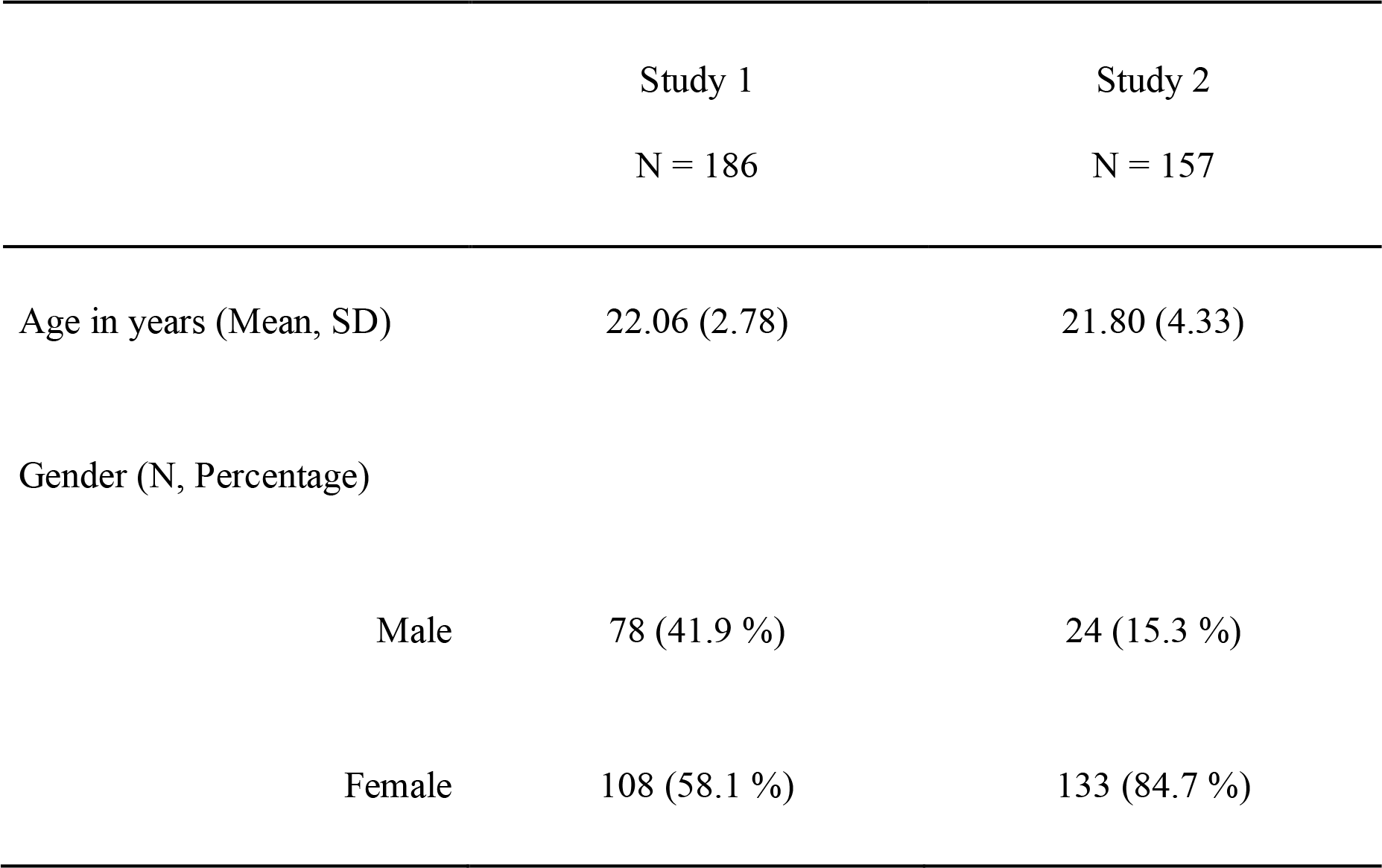
Descriptive statistics for the two samples.

### Tasks

#### Measure of statistical learning: Alternating serial reaction time (ASRT) task

In Study 1, implicit statistical learning was measured by a modified version of the ASRT task (Howard & Howard, 1997; Nemeth et al., 2010) (Figure 1a). In this task, participants were presented with a yellow arrow on the center of the screen pointing in one of four possible directions (left, up, down or right) for 200 ms. The presentation of the arrow was followed by a presentation of a fixation cross for 500 ms. Using a four-button Cedrus RB-530 response box, participants were instructed to press, as quickly as possible, the button corresponding to the direction of the arrow. Finger placement on each of the buttons of the response box was as follows: the up button had to be pressed with the left index finger; the down button had to be pressed with the right thumb; the right button had to be pressed with the right index; and the left button had to be pressed with left thumb. If participants responded correctly, the fixation cross would remain in the screen for another 750 ms. If participants did not answer or answered incorrectly, an exclamation mark or an “X” would appear for 500 ms, respectively, followed by a 250 ms fixation cross.

**Figure 1.**
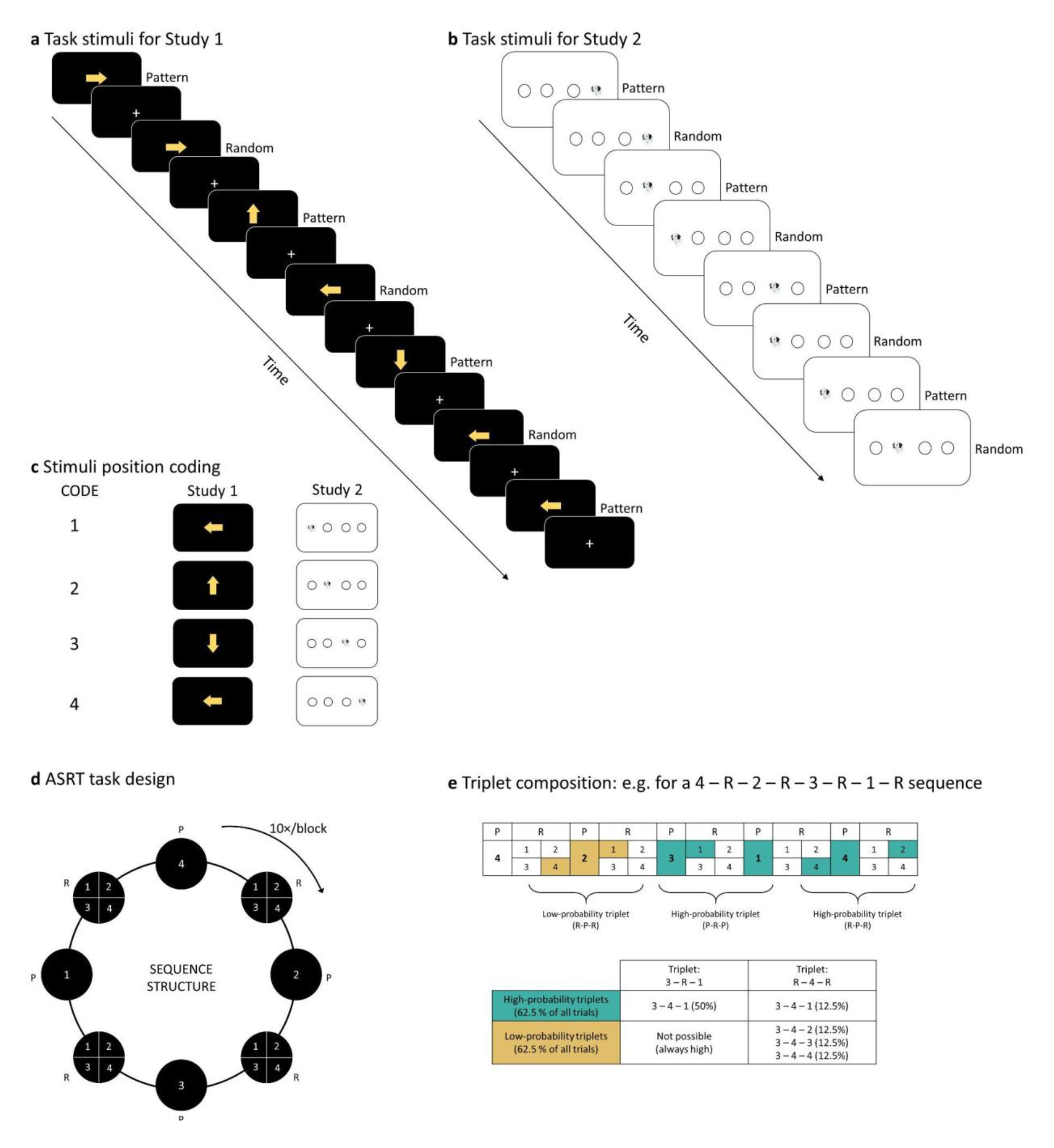
ASRT task design. **a)** In Study 1, the task stimuli consisted of yellow arrows that pointed in one of the four cardinal directions. A fixation cross was presented between each arrow. **b)** In Study 2, task stimuli were the head of a dog that appeared in one out of four different positions. **c)** Each stimuli position can be coded with a number. Here we have 1=left, 2=up, 3=down, 4=right for the arrows in Study 1, and 1=left, 2=center-left, 3=center-right, 4=right of the screen for the dog’s head in Study 2. **d)** The stimulus presentation followed an eight-element sequence, in which pattern (P) and random (R) stimuli alternated. The sequence was presented a total of 10 times per block. **e)** Sixty-four different triplets (runs of three consecutive stimuli) could result from the sequence structure. Some of the triplets appear more often than others. High-probability triplets could end in pattern or random element whereas low-probability triplets always ended with a random element. High-and low-probability triplets are denoted in green and yellow, respectively.

Unknowingly to the participants, the appearance of the stimuli followed a predetermined structure where pattern elements (P) alternated with random elements (R) (e.g. 2-R-4-R-3-R- 1-R, where the numbers represent the predetermined position of the stimuli, and “R” represent a random position) (Figure 1c &1d). Due to the pattern elements (P) alternating with random elements (R) in the ASRT task, some triplets (runs of three trials) had a higher probability of occurrence than others (Figure 1e). For example, in an 2-R-4-R-3-R-1-R sequence, the (2-3-4) or (3-4-1) triplets have a higher probability of occurrence compared to (2-3-2) or (4-3-1) triplets, as the former type of triplets can be found in P-R-P or in R-P-R structures whereas the second type of triplets can only occur in R-P-R structures. As a result, the former type of triplets are five times more likely to occur than the second type of triplets. Thus, the two types of triplets are referred to as high-and low-probability triplets, respectively. Previous studies using the ASRT task have consistently shown that, with increasing practice in the task, participants’ responses to the last element of high-probability triplets become faster compared to responses to the last element of low-probability triplets (Janacsek et al., 2012; Kóbor et al., 2017; Nemeth et al., 2010). Importantly, this phenomenon occurs without the emergence of an explicit knowledge of the sequence structure as reported by the participants (Janacsek et al., 2012; Kóbor et al., 2017; Vékony et al., 2022). Thus, implicit statistical learning in the task can be measured by computing the difference of reaction times between the last elements of high-probability triplets and the last element of low-probability triplets.

This first session consisted of 25 blocks of the ASRT task. There were 85 stimuli in each block, of which the first five were randomly ordered for practice purposes followed by 10 repetitions of the eight-element alternating sequence.

The ASRT task employed in Study 2 had four notable differences to the version used in Study 1. Firstly, in this version of the task, the stimuli (here, a drawing of a dog’s head instead of arrows) could appear in one of four horizontally arranged empty circles on the screen, instead of the four cardinal directions (Figure 1b). Secondly, the responses corresponded to the Z, C, B, and M keys on a QWERTY keyboard (with the rest of the keys removed). Thirdly, the task consisted of 45 blocks. We previously observed that although acceptable levels of reliability emerge even with 25 blocks, longer tasks lead to more reliable learning scores, which might be crucial for the correlational analyses we planned here (Farkas et al., 2022). Finally, the timing of the stimuli differed. The task in Study 2 was self-paced (i.e., the target stimulus remained on the screen until the correct response key was pressed) with a response-to-stimulus interval of 120 ms. Despite the differences, as detailed below, robust learning was observed in both studies, with similar distribution of learning scores, except an overall lower mean learning score in Study 1.

#### Neuropsychological tests for Study 1

##### Attentional network test (ANT)

The ANT allowed us to measure the capacities of three distinct networks of attention: the alerting, orienting and executive networks (Fan et al., 2002). This task required participants to determine, as fast as possible, whether a central arrow, presented in a set of five arrows, points to the left or to the right. The set of arrows can appear above or below a fixation cross and can be preceded or not by a spatial cue indicating their following location. Furthermore, the central arrow can be congruent (pointing in the same direction) or incongruent (pointing in a different direction) to the other arrows. We followed the standard calculation of network scores (Fan et al., 2002). The alerting component of attention was calculated by subtracting the mean RT of the central cue conditions from the mean of the no-cue conditions for each participant. In a similar manner, the orienting component of attention was calculated by subtracting the mean RT of the spatial cue conditions to the mean RT the center cue conditions and the executive component of attention was calculated by subtracting the mean RT of all congruent conditions to the mean RT of all incongruent conditions. In this task, higher scores for the alerting, orienting and executive scores indicate better attentional performances in these three aspects of attention.

##### Berg card sorting task (BCST)

We measured set shifting or cognitive flexibility using the computerised version of the BCST.64 available in the Psychology Experiment Building Language (PEBL) software (Mueller & Piper, 2014). In this test, a set of four cards on the top of the screen are presented to the participant. Each card has three characteristics: the colour, the shape and the number of items on the card. The participant was told to match new cards to the cards on the top of the screen according to one of the three characteristics, but they were not told which one; however, they received feedback about whether each choice they made was right or wrong. Thus, the participant was required to find the correct rule (matching characteristic) needed to match the cards as quickly and as accurately as possible. The participant was informed that the rule could change during the task. Cognitive flexibility in the BCST was measured by counting the perseverative errors, meaning the amount of errors reflecting lack of adaptation following a rule switch.

##### Counting Span (CSPAN) task

Updating or working memory capacity was measured by the CSPAN (Case et al., 1982). In this task, different shapes (blue circles, blue squares, and yellow circles) appeared on the computer screen. The participants’ task was to count out loud and retain the amount of blue circles (targets) among the other shapes (distractors) in a series of images. Each image included three to nine blue circles, one to nine blue squares and one to five yellow circles. At the end of each trial, participants repeated orally the total number of targets presented in the image and if counting was correct, the experimenter passed to the following trial When presented with a recall cue at the end of a set, participants had to rename the total number of targets of each image in their order of presentation. If the recall was correct, participants started a new set containing an extra image. The number of items presented in each image ranged from two to six. When participants made a mistake in the recall, the task was stopped and a new run, starting from a set with two trials, would start again. Each participant completed three runs of the task. Memory span capacity was computed as the mean of the highest set size the participant was able to recall correctly in the three runs.

##### Go No-go (GNG) task

We measured cognitive inhibition with the computerised version of the GNG task available in the Psychology Experiment Building Language (PEBL) software (Mueller & Piper, 2014). In the GNG task, participants were instructed to respond to certain stimuli (“go” stimuli) by clicking on a button as fast as possible and to refrain from clicking on other stimuli (“no-go” stimuli). In this version, participants were presented with a 2×2 array with four blue stars (one in the centre of each square of the array). Every 1500 ms, a stimulus (the letter P or R) would appear for 500 ms in the place of one of the blue stars. For the first half of the task, the letter P would be the “go” stimulus and the letter R would be the “no-go”. This rule would be then inverted in the second half of the task. Participants completed 320 trials. The ratio between “go” and “no-go” trials was 80:20, respectively. Cognitive inhibition capacity in the GNG task was measured by the d’:

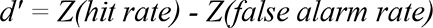

A higher d’ indicated better cognitive inhibition.

##### Verbal fluency tasks

Verbal fluency was measured with three subtasks testing the lexical, semantic and action components of verbal fluency. In the lexical fluency subtask, participants were required to say as many words as possible starting with the letter P. The letter P is often used in the French version of the phonemic fluency (St-Hilaire et al., 2016). In the semantic fluency subtask, participants were required to name animals, and in the action fluency subtask, isolated verbs describing an action realisable by a person. In each subtask, participants were instructed to say as many words as possible within one minute while avoiding word repetitions, words with the same etymological root and proper nouns. Every time the participant disrespected one of these rules, an error would be counted. Each verbal fluency subtask has a score computed by subtracting the amount of errors from the total amount of words produced within one minute. A higher score in each component of verbal fluency indicated a higher verbal fluency capacity.

#### Neuropsychological tests for Study 2

##### Counting Span (CSPAN) task

The procedure for measuring working memory capacity with the CSPAN task was identical in studies 1 and 2.

##### Digit Span (DSPAN) task

Phonological short-term memory capacity was measured using the DSPAN task (Isaacs & Vargha-Khadem, 1989). In this task, participants listened to and then repeated a list of digits enunciated by an experimenter. The task comprised seven levels of difficulty ranging from three to nine items. Each level comprised four different lists. If participants correctly recalled all items for at least three out of the four lists, they were permitted to move up a level. If participants could not recall at least three lists from the four lists correctly, the task ended. Memory span in the DSPAN task was considered to be the level (three to nine) at which the participant was still able to recall three from the four lists correctly. A higher memory span score is indicative of a better phonological short-term memory capacity.

##### Corsi blocks tapping task (Corsi) task

Visuo-spatial short-term memory capacity was assessed using the Corsi task (Kessels et al., 2000). In this task, nine cubes were placed in front of the participant in a fixed pseudo-random manner. The blocks were labelled with numbers only visible to the experimenter. The experimenter tapped a number of blocks in a specific sequence after which the participant had to tap the same blocks in the same order. Similarly to the DSPAN, the Corsi task comprised seven levels ranging from three to nine items. Four sequences were presented within a level. Memory span in the Corsi task was considered to be the level (three to nine) at which the participant was still able to recall three from the four lists correctly. A higher memory span score is indicative of a better visuo-spatial short-term memory capacity.

##### Verbal fluency tasks

Verbal fluency in Study 2 was measured with the Hungarian version of the task (Tánczos et al., 2014). Procedure was similar to the one used in Study 1 with two main differences: the lexical fluency was tested using the letter K and the action fluency was not measured in this study.

### Procedure

In Study 1 the experiment was organised over two sessions. During the first session, the ASRT task was administered. Participants were informed that the aim of the study was to study how an extended practice affected performance in a simple reaction time task. Therefore, participants were instructed to respond as fast and as accurately as they could. Participants were not given any information about the underlying structure of the task.

Participants completed the ASRT task in a soundproof experimental booth with a computer screen observable through a window and a four button (up, down, right left) response box placed over a table. Prior to the 25 blocks of the learning phase, the participants completed a three-block training phase where stimuli were completely random. Thus, the first three blocks contained no underlying sequence. This ensured that participants correctly understood the task instructions and familiarised themselves with the response keys. At the end of each block, participants received feedback on the screen reporting their accuracy and reaction time on the elapsed block. This was followed by a 15 second resting period. Participants were then free to choose when to start the next block.

The ANT, BCST, CSPAN, GNG and verbal fluency tasks were administered in the second experimental session. This session took approximately one hour. In order to avoid a possible fatigue effect in a particular task, the order of presentation of each task was randomised over participants.

The procedure for Study 2 was similar. The ASRT task was administered in the first session and CSPAN, DSPAN, Corsi and verbal fluency tasks were administered in a second session.

### Statistical analysis

#### ASRT learning trajectories

The ASRT blocks were compounded into units of five blocks (epochs) in order to facilitate data processing. ASRT task performance was assessed by calculating the median reaction times (RTs) of correct responses for high-and low-probability triplets separately in each epoch. We then computed learning scores for each epoch by subtracting the median RTs of high-probability triplets from the median RTs of low-probability triplets. A greater difference between high-and low-probability trials indicates greater learning. We excluded from the analysis trills (e.g., 2-1-2) and repetitions (e.g., 2-2-2) as participants show pre-existing tendencies to answer faster in these types of triplets. The first five trials (five warm-up random) of each block and trials with RTs below 100 ms were also removed from the analysis.

To evaluate SL, we conducted a repeated measures analyses of variance (ANOVA) with two factors: epoch (1-5) and triplet (high-vs. low-probability). Greenhouse-Geisser epsilon correction was used when Mauchly’s test of sphericity indicated that sphericity cannot be assumed. Original df values and corrected p values (if applicable) are reported together with partial eta-squared (η_p_^2^) as the measure of effect size. For the correlational analyses, a single index of implicit SL score for each participant was obtained by averaging the learning scores across epochs. Previous results from our group have indicated that while the reliability of learning scores is low for individual epochs, averaging across at least five epochs leads to acceptable levels of reliability. Thus, as adequate reliability is crucial for analyses of inter-individual differences, we decided to conduct our tests of the SL – EF relationship on averaged learning scores, instead of including executive functions score as a covariate in a learning trajectory analysis. We also complemented our classical analysis with Bayesian repeated measures ANOVA, conducted in JASP (JASP Team, 2019). Therefore, we additionally report the inclusion Bayes factors (BF) for each factor, obtained from Bayesian model averaging, and calculated from matched models only (see van den Bergh et al., 2020). The inclusion Bayes factor quantifies the change from prior inclusion odds to posterior inclusion odds and can be interpreted as the evidence in the data for including a predictor or factor. More generally, Bayes factors quantify the relative weight of evidence provided by the data for two theories, the null and the alternative hypotheses, H0 and H1 (Dienes, 2014). According to one commonly used classification (Wagenmakers et al., 2011), BF values between 1 and 3 indicate anecdotal evidence, values between 3 and 10 indicate substantial evidence, and values above 10 indicate strong evidence for H1. Conversely, values between 1/3 and 1 indicate anecdotal, values between 1/10 and 1/3 indicate substantial evidence, and values below 1/10 indicate strong evidence for H0. Values close to 1 do not support either H0 or H1.

#### Factor analyses

We further aimed to determine the potential latent structure of our sets of EF measures in a data-driven manner. Therefore, we investigated the factor structure of the EF measures in both of our datasets separately, using maximum likelihood exploratory factor analysis (ML EFA) with varimax rotation, as implemented in the psych package in R (Revelle, 2022), with default settings. To aid interpretation, the scores that reflect error percentages were subtracted from 100, so that higher values represent better performance in all variables. To assess the factorability of the data, we utilised 3 complementary approaches (Dziuban & Shirkey, 1974). Firstly, we inspected the off-diagonal elements of the anti-image covariance matrix. If the dataset is appropriate for factor analysis, these elements should be all above .50. Secondly, we computed the Kaiser-Meyer-Olkin (KMO) test of sampling adequacy. The higher the overall KMO index, the more appropriate a factor analytic model is for the data. The original cut-off recommended by (Kaiser, 1970) is .60, but other authors have also suggested .50 (Dziuban & Shirkey, 1974). Finally, we performed Bartlett’s test of sphericity, which tests the hypothesis that the sample correlation matrix came from a multivariate normal population in which the variables of interest are independent (Bartlett, 1950). Rejection of the hypothesis is taken as an indication that the data are appropriate for analysis.

We determined the number of factors to extract using Horn’s parallel analysis (Horn, 1965). This approach is based on comparing the eigenvalues of factors of the observed data with those of random data from a matrix of the same size. Factors with higher eigenvalues in the observed, than in the random data are kept. We used a more stringent criteria, and compared the observed eigenvalues to the 95th percentile, instead of the mean of the simulated distributions. Furthermore, our use of ML EFA also allowed us to calculate multiple fit indices of the applied factor analytic models. Following the recommendations of Fabrigar et al. (1999) and Hu & Bentler (1998), we chose to focus on the Root Mean Square Error of Approximation (RMSEA) and the Standardized Root Mean Squared Residual (SRMR). According to a commonly used guideline, RMSEA values less than 0.05 constitute good fit, values in the 0.05 to 0.08 range acceptable fit, values in the 0.08 to 0.10 range marginal fit, and values greater than 0.10 poor fit. Similarly, SRMR values below .08 are generally considered indicators of good model fit (Hu & Bentler, 1999).

After we determined the number of factors to extract, participant level factor scores were calculated for all factors, based on Thomson’s (1939) regression method. Correlations between these scores and the ASRT were then assessed. For these correlations, we also calculated Bayes factors, using JASP (JASP Team, 2019), with default priors (stretched beta distribution with a width of 1). We tested the alternative hypothesis that the two variables are negatively correlated. We also tested the robustness of our Bayes factors to different prior widths.

#### Meta-analysis

In addition to testing our hypotheses in each of the two samples separately, following the recommendation of Braver et al. (2014), we also carried out a continuously cumulating meta-analysis (CCMA) of the two studies. A meta-analytic approach allows us to pool individual effects sizes into a single estimate, while quantifying their heterogeneity. In order to make the two studies comparable, in the two datasets separately, we extracted factor scores from a single factor ML EFA applied to the set of EF tasks that were shared in both studies. These were Lexical fluency, Semantic fluency and CSPAN. We then calculated the Pearson’s correlation between these EF factor scores and the implicit SL learning scores and ran our meta-analytic model on these effects. We fit a fixed-effect meta-analysis model implemented in the ‘meta’ R package (Harrer et al., 2021; Schwarzer, 2022), using the inverse variance pooling method. We expected little between-study heterogeneity in the effect, therefore a priori, we decided on a fixed-effect model to test the average true effect in the two studies. We nevertheless estimated between-study heterogeneity using the REML estimator (Viechtbauer, 2005). As reported below, commonly used measures of heterogeneity indicated little between-studies variability, validating our choice of a fixed-effect model. We relied on the REML estimate of the variance of the distribution of true effects, τ2, on the percentage of variability in the effect sizes that is not caused by sampling error, I2 (Higgins & Thompson, 2002), and on Cochran’s Q (Cochran, 1954), which can be used to test whether there is more variation than can be expected from sampling error alone.

## Results

Descriptive statistics of the samples are presented in Table 1. Histograms showing the distributions of all variables are presented in Supplementary Figure S2 and S3, for Study 1 and 2, respectively.

### Implicit statistical learning

#### Study 1

To investigate implicit SL in the ASRT task, we analysed median RTs by a repeated measures ANOVA with two factors: EPOCH (1-5) and TRIPLET (high-vs. low-probability). The main effect of EPOCH was significant (*F*_(4, 740)_ = 27.264, *p* < .001, η_p_^2^ = .128, BF = 1.154e+18). indicating general skill learning, i.e., participants became faster overall as the epochs went on, irrespective of triplet probabilities (Figure 2a). The main effect of TRIPLET was significant (*F*_(1, 185)_ = 87.855, *p* < .001, η_p_^2^ = .322, BF_inclusion_ = 7.013e+13), with average responses to high-probability triplets being faster than responses to low-probability triplets, revealing successful SL. The EPOCH x TRIPLET interaction was also significant (*F*_(4, 740)_ = 5.027, *p* < .001, η_p_^2^ = .026, BF_inclusion_ = 12.291). Post-hoc tests revealed significantly faster responses to high-as compared to low-probability triplets in all epochs (all p_bonf_ < .012), except in epoch 2 (p_bonf_ = .084), with progressively larger effect sizes, indicating that SL improved over time as the task went on (also see Figure 2a).

**Figure 2.**
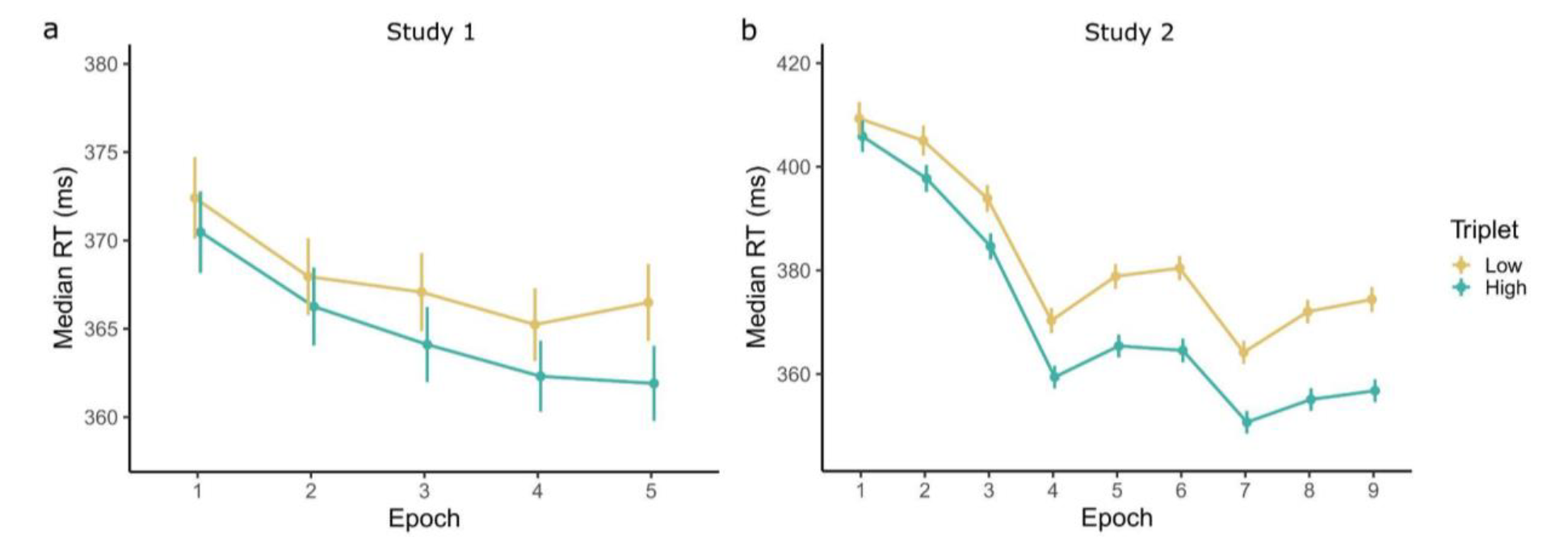
Epoch-wise median reaction times (RTs) in the ASRT tasks for high-and low-probability triplets. **a)** In Study 1 and **b)** in Study 2. There was a significant RT difference between high and low triplets showing statistical learning. SL improved significantly over time. Error bars indicate standard error of the mean. Note the different scales of the y axis.

#### Study 2

We carried out the same repeated measures ANOVA for Study 2 (the sole difference being that the EPOCH factor now had 9 levels), and found similar learning trajectories (Figure 2b). The main effect of EPOCH was significant (*F*_(8, 1248)_ = 372.518, *p* < .001, η_p_^2^ = .705, BF = ∞) indicating general skill learning, i.e., participants became faster overall as the epochs went on, irrespective of triplet probabilities. The main effect of TRIPLET was significant (*F*_(1, 156)_ = 540.716, *p* < .001, η_p_^2^ = .776, BF = 7.386e+48), with average responses to high-probability triplets being faster than responses to low-probability triplets, revealing successful SL. The EPOCH × TRIPLET interaction was also significant (*F*_(8, 1248)_ = 35.405, *p* < .001, η_p_^2^ = .185, BF_inclusion_ = 1.989e+4). Post-hoc tests revealed significantly faster responses to high-as compared to low-probability triplets in all epochs (all p_bonf_ < .026), with progressively larger effect sizes, indicating that SL improved with practice as the task went on (also see Figure 2b).

### Correlations between procedural learning and executive functions

#### Study 1

Bivariate Pearson’s correlations between all variables are presented in Figure 3a. EF measures tended to be weakly or moderately positively correlated, the strongest relationships seemed to be between the fluency measures, and between the fluency measures and CSPAN.

**Figure 3.**
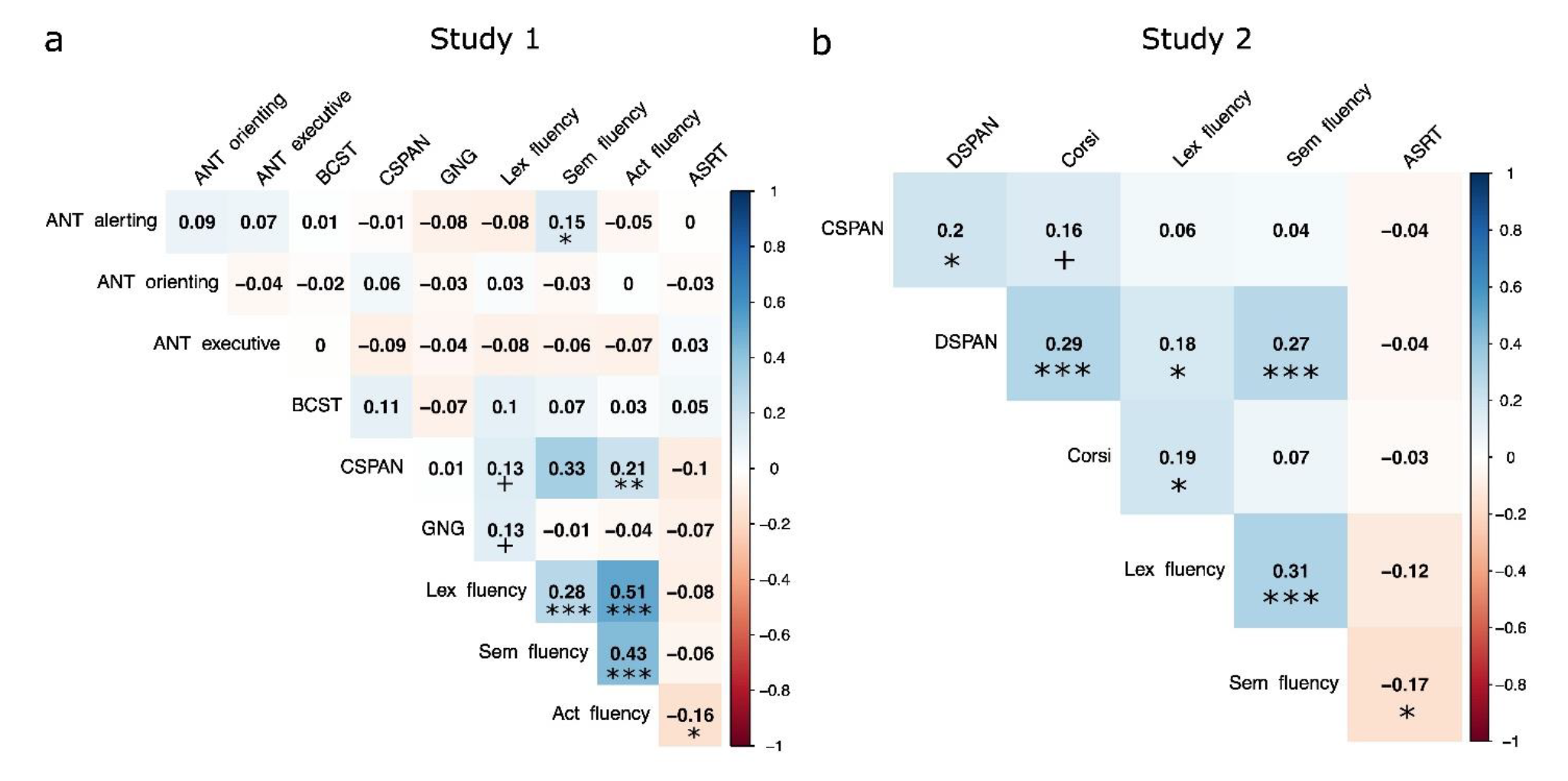
Bivariate correlations between the EF measures and the ASRT. **a)** In Study 1 and **b)** in Study 2. More negative and more positive correlations are indicated by red and blue backgrounds, respectively. The working memory and verbal fluency measures correlated moderately positively with each other, and weakly negatively with ASRT. ***p<.001, **p<.01, *p<.05, +p<.10

Implicit SL ability had generally negative correlations with EF measures, the strongest of these was with Action fluency (Pearson’s r = -.165, 95% CI = [-.302, -.021], *p* = .025, Speaman’s rho = -.162, 95% CI = [-.299, -.018], *p* = .027).

#### Study 2

Bivariate Pearson’s correlations between all variables of Study 2 are presented in Figure 3b. EF measures tended to be weakly or moderately positively correlated, again the fluency measures correlated the strongest.

Implicit SL ability had generally negative correlations with EF measures, the strongest of these was with Semantic fluency (Pearson’s r = -.167, 95% CI = [-.315, -.011], *p* = .037, Spearman’s rho = -.145, 95% CI = [-.295, .012], *p* = .070).

### Factor analyses of executive function measures

Besides investigating the relationship between performance on individual EF tasks and implicit SL, we also aimed to investigate the relationship between implicit SL and general EF ability that might be captured by shared variance on all tasks. To extract such a common EF measure from the individual tasks, we used Maximum Likelihood Exploratory Factor Analysis (ML EFA). We opted for EFA in order to find a set of latent constructs that capture our specific set of EF measures, while remaining a priori agnostic about the exact structure of these components. While the ML approach allows the computation of various goodness of fit indices, that are unavailable to principal factor approaches.

#### Study 1

The dataset we considered for factor analysis consisted of the set of EF measures presented in Figure 3a. Beforehand, we established the factorability of the data using 3 approaches. Firstly, the diagonals of the anti-image correlation matrix of the data were all over .5, which suggests good factorability. Secondly, the overall Kaiser-Meyer-Olkin (KMO) measure of sampling adequacy was .59, which is somewhat below the cut-off value of .6 originally suggested by Kaiser (1970), but above the cut-off of .5, suggested by other authors (Dziuban & Shirkey, 1974). Finally, Bartlett’s test of sphericity was significant (χ2 (36) = 145.029, *p* < .001), suggesting adequate factorability. Based on this, the data were deemed appropriate for factor analysis.

To determine the number of factors to extract, we relied on parallel analysis, and goodness of fit indices. Parallel analysis suggested that 1 factor was extractable, based on the comparison of the eigenvalues with randomly generated data (Figure 4a). Thus, the single factor solution was selected.

**Figure 4.**
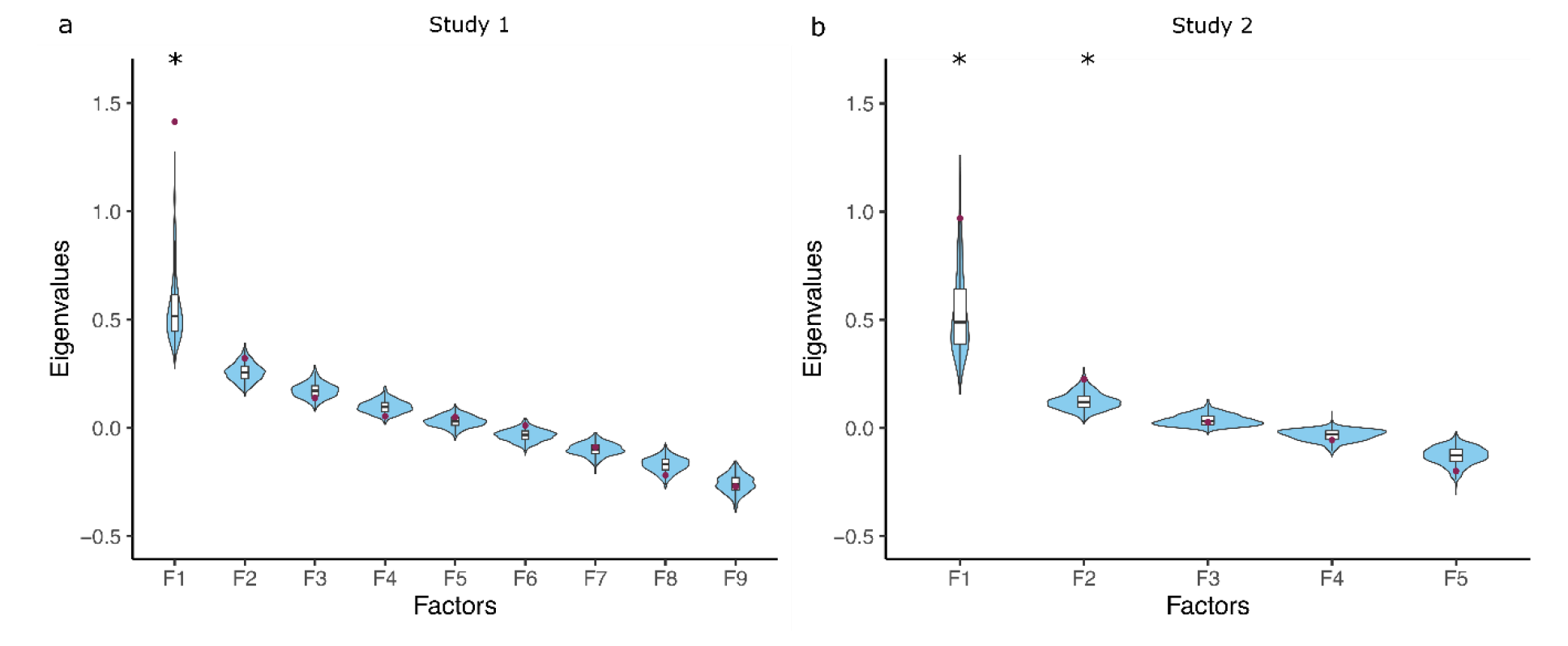
Parallel analysis to determine the extractable factors. **a)** In Study 1 and **b**) in Study 2. The observed eigenvalues are indicated by red dots. These were compared to the distribution of eigenvalues obtained from simulated data, indicated by violin plots and boxplots. Stars indicate factors for which the observed eigenvalue is larger than the 95th percentile of the simulated distributions, and thus were deemed extractable.

This factor explained 16% of the variance. The χ2 test of the null hypothesis that one factor is sufficient was not significant (χ2 (27) = 36.87, *p* = .098), suggesting that the null can be accepted, meaning that our single factor was sufficient in capturing the full dimensionality of the data. Goodness of fit indices were also indicative of good fit (RMSEA = .044, 95% CI = [.000 .077]; SRMR = .062). Factor loadings are presented in Table 2. Factor 1 had high loadings from the three fluency variables and from CSPAN, confirming the pattern of bivariate correlations. We calculated factor scores for each subject, and investigated the correlations with implicit SL.

**Table 2.**
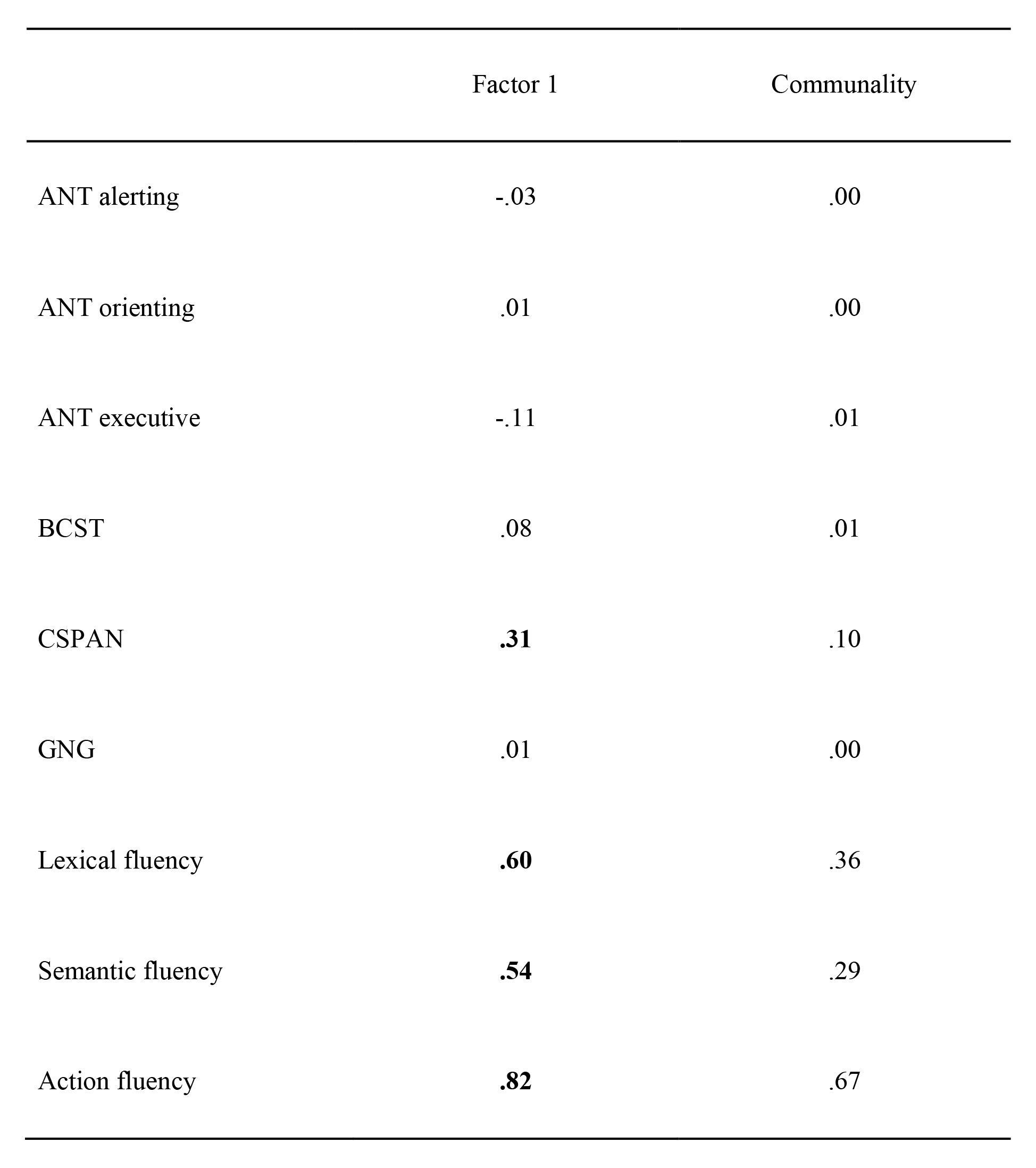
Study 1. Factor loadings and communalities of each EF measure on the factors, based on the 1 factor ML solution (16% of variance explained). Loadings above the threshold of .30 are highlighted in bold to aid interpretation.

Factor 1 significantly negatively correlated with implicit SL (Figure 5a, Pearson’s r = -.156, 95% CI = [-.293, -.012], *p* = .034, Spearman’s rho = - .154, 95% CI = [-.292, -.011], *p* = .036). The Bayes factor for the one-sided alternative hypothesis that the two variables are negatively correlated was BF-0 = 1.683, meaning that the data are approximately 1.7 times more likely to occur under the alternative hypothesis, indicating anecdotal evidence in favour of it. Robustness checks of the Bayes factor to varying prior distribution width are included in the .jasp files in Supplementary Materials. Histograms showing the distribution of the factor score, as well as the original EF measures are similarly presented in the supplementary material.

**Figure 5.**
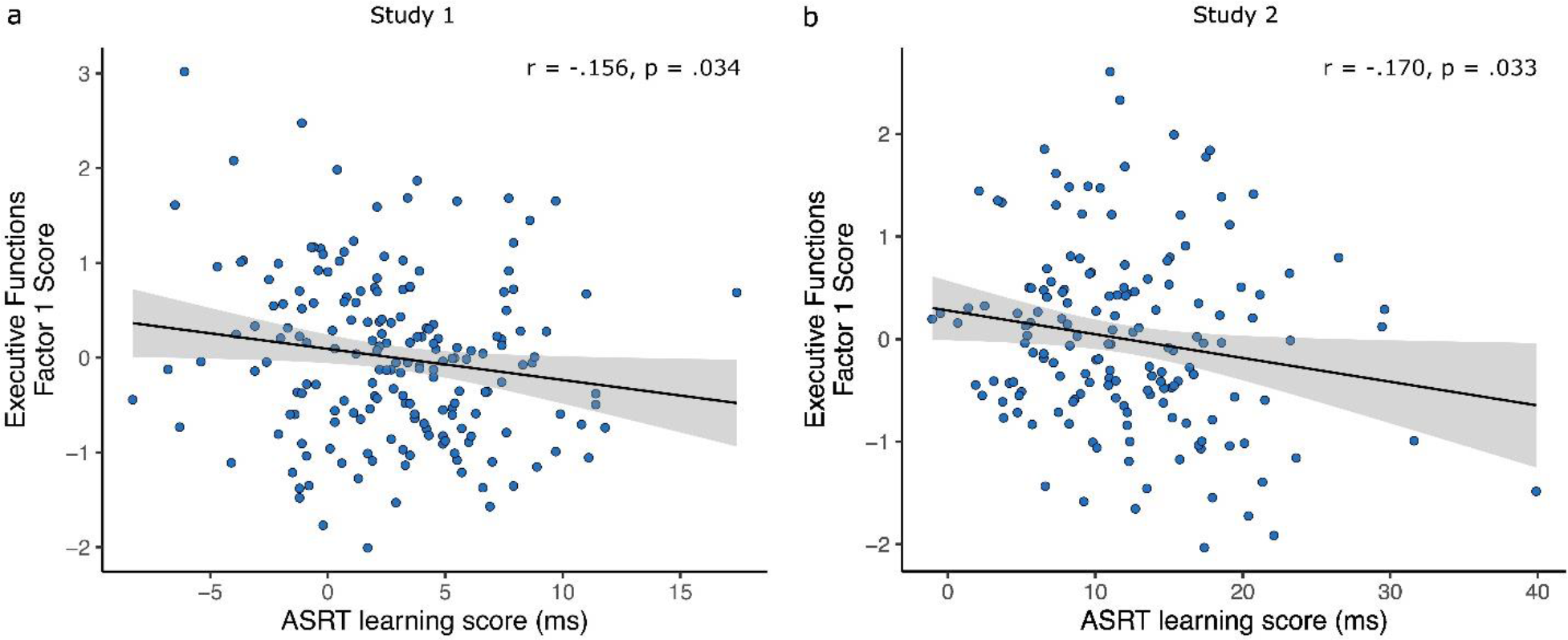
Relationship between ASRT learning scores and EF Factor 1 scores across subjects. Solid line is the linear fit, shaded area corresponds to the 95% CI. One datapoint per participant. **a)** In Study 1 and **b**) in Study 2.

#### Study 2

We proceeded in the same manner for Study 2. The dataset we considered for factor analysis consisted of the set of EF measures presented in Figure 3b. We followed the same approach to establishing the factorability of data, as in the analysis of Study 1. The diagonals of the anti-image correlation matrix of the data were all over .5. The overall KMO measure of sampling adequacy was .60, meeting both more conservative and more liberal cut-off criteria. Finally, Bartlett’s test of sphericity was significant (χ2 (10) = 54.591, *p* < .001). Based on this, the data were deemed appropriate for factor analysis.

To determine the number of factors to extract, we again relied on parallel analysis. Parallel analysis suggested that 2 factors are extractable, based on the comparison of their eigenvalues with random data (Figure 4b). Thus, the 2 factor solution was selected.

These two factors explained 19% and 14% of the variance, respectively. The total variance explained of the model was thus 34%, somewhat more than the explained variance of the Study 1 EFA model. The χ2 test of the null hypothesis that 2 factors are sufficient was not significant (χ2 (1) = 1.40, *p* = .236), suggesting that the null can be accepted, meaning that our 2 factors are sufficient in capturing the full dimensionality of the data. Goodness of fit indices were also indicative of good fit (RMSEA = .050, 95% CI = [.000 .227]; SRMR = .020), although the RMSEA had a noticeably larger CI, compared to the Study 1 EFA model. Factor loadings are presented in Table 3. Factor 1 had high loadings from the 2 fluency variables and a somewhat weaker loading from DSPAN. Factor 2 had high positive loadings from the CSPAN, DSPAN and Corsi tasks. Thus, the factor structure reflected the dissociation between the fluency and the short-term memory measures.

**Table 3.** Study 2. Factor loadings and communalities of each EF measure on the 2 factors, based on the 2 factor varimax rotated ML solution (34% of variance explained). Loadings above the threshold of .30 are highlighted in bold to aid interpretation.

|  | Factor 1 | Factor 2 | Communality |
| --- | --- | --- | --- |
| CSPAN | .03 | **.32** | .10 |
| DSPAN | .28 | **.52** | .34 |
| Corsi | .08 | **.54** | .30 |
| Lexical fluency | **.33** | .23 | .16 |
| Semantic fluency | **.88** | .03 | .77 |

Factor 1 significantly negatively correlated with implicit SL, and with a similar effect size, as in Study 1 (Figure 5b, Pearson’s r = -.170, 95% CI = [-.319, -.014], *p* = .033, Spearman’s rho = -.150, 95% CI = [-.300, .007], *p* = .060). The Bayes factor for the one-sided alternative hypothesis that the two variables are negatively correlated was BF-0 = 1.875, meaning that the data are approximately 1.9 times more likely to occur under the alternative hypothesis, indicating anecdotal evidence in favour of it. Factor 2 did not correlate significantly with implicit SL ability (Factor 2: Pearson’s r = -.034, 95% CI = [-.190, .123], *p* = .672, Spearman’s rho = -.023, 95% CI = [-.179, .134], *p* = .771, BF-0 = 0.145). Robustness checks of the Bayes factor to varying prior distribution width are included in the .jasp files in Supplementary Materials. Histograms showing the distribution of the 2 factor scores, as well as the original EF measures are similarly presented in the supplementary material.

### Continuously Cumulating Meta-Analysis

Besides testing the hypothesis of a negative association between common EF ability and implicit SL in our two samples separately, we also ran a fixed-effect meta-analytic model, in order to pool evidence from both of our studies into a single effect size, while estimating their heterogeneity. Heterogeneity metrics revealed little between-studies variability in the effect, validating our choice of a fixed-effect model. The between-study heterogeneity variance was estimated at τ^2^ = 0, with an I^2^ value of 0%, Cochran’s Q test was also not statistically significant, Q(1) = 0.60, p = .440. The pooled effect size was negative and significantly different from zero, r = -.115, 95% CI = [-.218, -.008], *p* = .035 (Figure 6). The weights of Study 1 and Study 2 in the pooled effect were 54.3% and 45.7%, respectively. We also carried out this analysis using factor scores derived from all EF measures in both tasks, instead of just the shared ones (Supplementary Figure S1). This analysis also yielded a pooled effect size, significantly more negative than 0, r = -.162, 95% CI = [-.264, -.057], *p* = .003.

**Figure 6.**
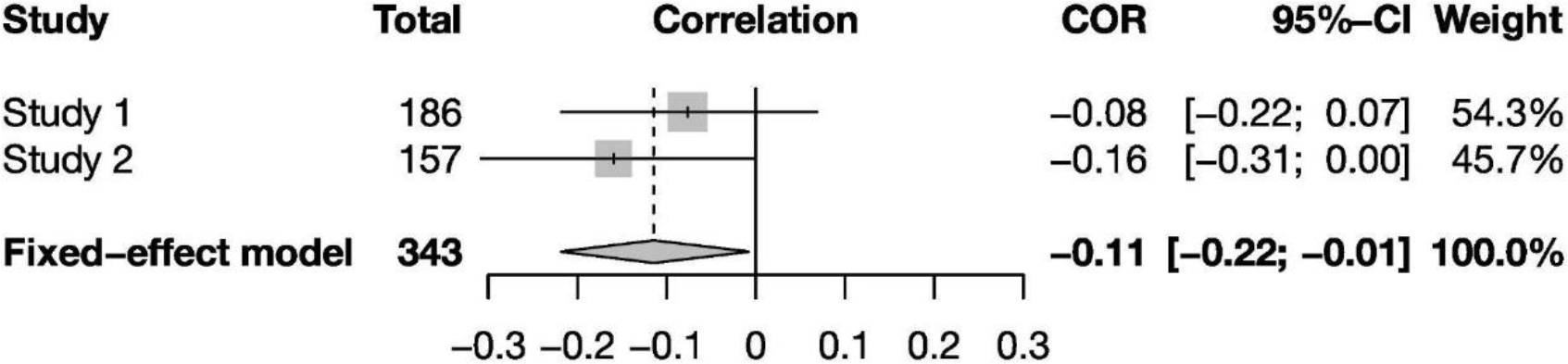
CCMA of the two studies. Pearson’s r between EF factor scores and implicit SL learning scores, along with its 95% CI is shown next to each individual study. Below, the pooled effect size from a fixed-effect meta-analytic model and its 95% CI is shown. Individual and total sample sizes are also indicated, as well as study weight. Factor scores were derived only from tasks that were shared by both studies.

## Discussion

In this study, we explored how implicit SL relates to multiple components of prefrontal lobe-dependent EF. At the level of individual tasks, SL correlated negatively with most EF measures, the strongest of these correlations being with verbal fluency. At the level of latent EF components, an EF factor comprising verbal fluency and counting span scores in Study 1 and verbal fluency and digit span scores in Study 2 also correlated negatively with SL, with a modest, but similar effect size in both studies. Our results imply that individuals with better verbal fluency and working memory ability have lower implicit statistical learning ability, suggesting that specific prefrontal lobe functions might interfere with implicit statistical learning while still allowing for the acquisition of the underlying regularities. We aimed to improve upon previous studies and extend their results in several ways. We made use of far larger samples (N_Study1_ = 186, N_Study2_ = 157), giving us larger statistical power than any previous study in the field (Button et al., 2013). Our successful internal replication, using a second sample and a cumulative meta-analytic approach, speaks to the robustness of our results (Braver et al., 2014). Our measurement of EF components at both the task and the latent variable level using exploratory factor analysis allowed for a more thorough characterization of this complex cognitive function (Diamond, 2013), and attenuated measurement error (Haines et al., 2023), while remaining agnostic about the measurement model of EF (Karr et al., 2018). Finally, we used an implicit SL task that has been shown to be both valid (Buffington et al., 2021) and reliable (Farkas et al., 2023), which is crucial, given the recent debate about the psychometric properties of commonly used implicit learning tasks (West et al., 2018).

We interpret our results in the PFC-mediated competition model of declarative-procedural interactions (Freedberg et al., 2020; Janacsek & Nemeth, 2022; Poldrack & Packard, 2003), and suggest that the observed negative relationship between EF and SL might be due to the suppression of model-free, procedural learning by prefrontal EF. Support for this theory has come not only from the effects of PFC disruption on learning (Ambrus et al., 2020; Borragán et al., 2016; Filoteo et al., 2010; Nemeth et al., 2013; Smalle et al., 2017, 2022) and from computational modelling (Lee et al., 2014) studies, but also from neural data. For example, a meta-analysis of multiple primate studies by Loonis et al. (2017) has revealed distinct patterns of post-choice oscillatory synchrony within PFC during implicit versus explicit learning, such that Delta/Theta band synchrony increased after correct choices during implicit learning, but after incorrect choices during explicit learning. Moreover, their results also suggested that whereas explicit learning was associated with increased synchrony between PFC and hippocampus in the alpha and beta bands, implicit learning was associated with decreased synchrony between PFC and caudate in the theta band. A similar pattern of results was revealed in humans by Voss et al. (2012), who showed that the use of a flexible, declarative learning strategy was linked to the interaction between the medial temporal lobe and the fronto-parietal attentional network, whereas the use of a more rigid, procedural learning strategy was linked to caudate nucleus fronto-parietal network interactions. Furthermore, neuroimaging studies also indicate that statistical learning is associated with decreased functional connectivity within PFC circuits and between the PFC and other networks (Park et al., 2022; Tóth et al., 2017).

In Study 1, the most parsimonious model consisted of a single factor, comprising fluency and counting span performances. The link between verbal fluency and working memory tasks is not surprising. Fluency scores have constantly been found to relate to verbal working memory (Daneman, 1991; Rende et al., 2002; Unsworth et al., 2011; Unsworth & Engle, 2005; Woods et al., 2005). Higher working memory capacity may aid in handling the cognitive load in fluency tasks (Rosen & Engle, 1997), while lower capacity may lead to more perseveration errors (Azuma, 2004). Shao et al. (2014) have also found that both category and letter fluency scores were uniquely predicted by updating in working memory. Similar findings that relate updating to verbal fluency performance were found by Gustavon et al. (2019). The factor structure uncovered by Fisk & Sharp (2004) is also suggestive of this relationship. Although they loaded onto a separate factor more strongly, a factor representing updating also had quite high loadings from word fluency tasks. Thus, the factor in Study 1 may reflect updating. This makes sense when considering the demands of verbal fluency and counting span tasks. Both tasks require tracking and updating sequences of items. In verbal fluency tasks, individuals generate words fitting a specific category, requiring constant updating of working memory. Without updating, they may repeat words or struggle to generate new ones, affecting performance. Counting span tasks involve maintaining and updating a number sequence while performing a secondary task. Without updating, individuals may lose track of the sequence, negatively impacting performance. Updating appears to be a suitable explanation for the common factor in verbal fluency and counting span tasks. Although inhibition and set shifting have been proposed as significant factors in fluency tasks (Shao et al., 2014), the absence of positive correlations between our measures of shifting (BCST) and inhibition (GNG) and other executive function tasks makes it challenging to consider them as appropriate explanations for factor 1.

While our general findings, suggestive of a negative relationship across individuals in EF and SL are in line with the competitive neurocognitive systems framework (Janacsek & Nemeth, 2022; Poldrack & Packard, 2003; Reber, 2013), the fact that it is primarily for the updating component that this association was uncovered runs contrary to some previous results showing a positive relationship between working memory and statistical learning (Frensch & Miner, 1994; Howard & Howard, 1997; Park et al., 2020). The central idea of these studies was that larger working memory capacity might open up a larger “window” for serial order learning. However, as noted by Janacsek and Nemeth (2013), these effects seem primarily observed in explicit learning conditions and consolidation, rather than implicit learning of probabilistic representations per se. Our results imply that under implicit learning, EF updating might instead compete with statistical learning. A possible explanation might be that updating might disrupt the stabilization of a simple predictive model allowing triplet learning in the ASRT task. However, we note that the nature of this kind of “updating” of implicit probabilistic representations is likely quite different from the explicit updating of items in working memory that is involved in EF tasks. Cognitive control, executive functions and working memory are related to a model-based computational strategy (Otto, Gershman, et al., 2013; Otto, Raio, et al., 2013). So our results might be interpretable as showing that more automatic, model-free learning is antagonistically related to model-based updating associated with goal-directed processes.

An alternative explanation to updating is that both working memory and fluency are strongly related to long-term memory access. Numerous theories and empirical evidence support the former (Cowan, 2016; Unsworth et al., 2013), while the fluency test merely involves retrieving words from the mental lexicon (Ullman, 2007). From this, we can speculate that if access to our long-term representations and models is poorer, it significantly enhances model-free learning because there is no interference from previous models in learning new patterns and predictions. This explanation aligns with the explanation provided by Ambrus et al. (2020), who attributed their results of inhibitory TMS-enhanced statistical learning to difficulties in long-term memory access or the suppression of top-down processes. However, this hypothesis needs to be validated through studies that examine long-term memory access, cognitive control, and predictive processes not only through behavioral but also neuroimaging techniques within a single experimental design.

While our study has notable strengths, including the incorporation of two large, independent samples, multiple EF measures, and the use of data-driven latent component extraction, there are some limitations that need to be mentioned, and addressed in future research. Firstly, the effects sizes were relatively small. This likely stems partially from the wide array of factors that influence EF and SL ability, and partially from attenuated correlations due to measurement error. We have recently estimated the reliability of the ASRT task in similar settings as the current study, to be between .754 and .791. If we correct our correlation coefficients for attenuation with the lowest reliability value from that study, we obtain 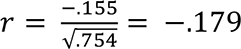 and 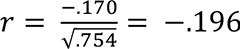, which are likely better estimates of the true relationships. Secondly, to replicate the results obtained in Study 1, we re-analysed data from a sample previously collected by us, and first described in Kóbor et al. (2017). While we note that as this is an earlier study, which was not designed with the research question of this study in mind, we believe the differences between protocols do not meaningfully diminish the informativeness of our replication attempt. Indeed, the two primary differences are the different set of EF tasks, and the longer ASRT. Regarding the first, the results of Study 1 show that tasks tapping into the updating component of EF (CSPAN and verbal fluency) showed the clearest relationship with implicit SL. These are exactly the tasks that were shared between studies. Regarding the second study, we have previously shown that longer ASRT tasks lead to more reliable learning scores (Farkas et al., 2022), which are crucial for correlational research designs. Thus, if anything the longer task should increase the power of our replication attempt. However, the different response to stimulus intervals of the two studies does decrease their comparability. As a result, it would be beneficial for future studies and replication attempts to show a greater consistency across studies in the protocol. Finally and relatedly, the incomplete nature of our EF battery should also be addressed by future work. Our set of EF tasks did not cover the most widely accepted theoretical model of EF components in an equal manner, as for set shifting and inhibition, we only had the BCST and GNG tasks, respectively (Miyake et al., 2000). Thus, our results should be extended and confirmed by future studies with a more complete set of EF tasks, and a more theory-driven assessment of latent EF components, for example with Confirmatory Factor Analytic models. We note however, that the ubiquity of the Unity and Diversity model has been questioned recently (Karr et al., 2018).

To our knowledge this was the first study to combine multiple data sets to investigate the relationship between implicit SL and EF testing large pools of participants. We found that general executive function has a negative relationship with implicit SL, suggesting that individuals with better EF functions might be worse at acquiring probabilistic models of visuospatial information. Our results fall in line with the theoretical framework stating that implicit automatic processing works in competition with PFC functions. This study takes a step further by exploring which components of executive functioning might be at the core of this competition. Our results are indicative that it is the updating component that is driving this relationship. Our study highlights the importance of exploring the relationship between different cognitive abilities using larger sets of subjects and in a data-driven manner.

### Credit author statement

Felipe Pedraza: Conceptualization, Methodology, Validation, Formal analysis, Investigation, Data curation, Writing – Original draft, Writing – Review & Editing, Visualization

Bence C. Farkas: Conceptualization, Methodology, Software, Validation, Formal analysis, Investigation, Data curation, Writing – Original draft, Writing – Review & Editing, Visualization

Teodóra Vékony: Conceptualization, Methodology, Software, Validation, Formal analysis, Investigation, Data curation, Writing – Review & Editing, Visualization, Supervision, Project administration

Frederic Haesebaert: Inclusion of participants, Writing – Review & Editing

Karolina Janacsek: Conceptualization, Methodology, Investigation, Data curation, Project administration, Funding acquisition

Royce Anders: Writing – Review & Editing

Barbara Tillmann: Writing – Review & Editing

Gaën Plancher: Writing – Review & Editing

Dezso Nemeth: Conceptualization, Methodology, Investigation, Data curation, Writing – Review & Editing, Supervision, Project administration, Funding acquisition

### Open practices

All code and data necessary to replicate the findings of this manuscript can be found on the OSF platform, at the following link: https://osf.io/2asnb/

## Acknowledgements

The authors are grateful to Jeoffrey Maillard for managing participant enrollment. This work was supported by the Chaire de Professeur Junior Program by INSERM and French National Grant Agency (Chaire Inserm-ANR-22-CE64-001); the National Brain Research Program (NAP2022-I-2/2022), János Bolyai Research Scholarship of the Hungarian Academy of Sciences (to K.J.). Project no. 124148 has been implemented with the support provided by the Ministry of Innovation and Technology of Hungary from the National Research, Development, and Innovation Fund, financed under the NKFI/OTKA PD funding scheme (to K.J.).

